# CAR T cells for solid tumors: genome-wide dysfunction signature confirms value of c-Jun overexpression, but signals heterogeneity

**DOI:** 10.1101/2020.02.07.939033

**Authors:** Mostapha Benhenda

**Affiliations:** Melwy

## Abstract

Chimeric antigen receptor (CAR) T cells still have limited effects in cancer, and especially in solid tumors, due to T cell dysfunction and exhaustion. CAR T cells overexpressing c-Jun (JUN CAR T cells) have been introduced to solve this problem. In this paper, we analyze JUN CAR T cells scRNA-seq data in solid tumors, by applying a genome-wide signature of T cell dysfunction, TID. This signature comes from the bulk RNA-seq signature TIDE, introduced to predict immune checkpoint inhibitor response. Our analysis confirms that on average, JUN CAR T cells are less dysfunctional than non-JUN CAR T cells. However, it also shows heterogeneity within JUN CAR T cells, which brings uncertainty about possible tumor resistance. We conclude that genome-wide dysfunction signature TID helps de-risking CAR T cell therapy for solid tumors.

## Introduction

CAR T cells still have limited results against solid tumors, despite their successes in some hematological malignancies. One reason of failure is T cell dysfunction, due to T cell exhaustion [2, 7, 8]. To address this issue, [5] modifies CAR T cells of the HER2-BBz type, in order to overexpress c-Jun, an AP-1 transcription factor associated with productive T cell activation.

In mouse models of osteosarcoma (a solid tumor affecting bones), they report better survival [5, extended data figure 9h]. Moreover, they report less exhaustion, and better activation for JUN HER2-BBz CAR T, compared with control HER2-BBz CAR T cells. For the most part, they rely on a single-cell analysis of single-gene expression signatures, such as PDCD1, IL2 and CD28.

However, in solid tumor transcriptomics, single-gene signatures are now not sufficient. They are complemented with genome-wide signatures, which, for example, better predict response to immune checkpoint inhibitors than single-gene signatures like PDL1 [4].

Among all genome-wide signatures, Tumor Immune Dysfunction and Exclusion, or TIDE [4], is particularly relevant, because it includes a distinctive T cell dysfunction signature, TID (Tumor Immune Dysfunction). In this paper, we apply TID signature to CAR T single-cell RNA-seq data from [5].

### Single-gene signatures

In [5], CAR T cell data is analyzed using several single-gene signatures. Among them, we consider the 12 signatures that appear in [5, figure 6g] (copied here in figure 1a):

**Figure 1:**
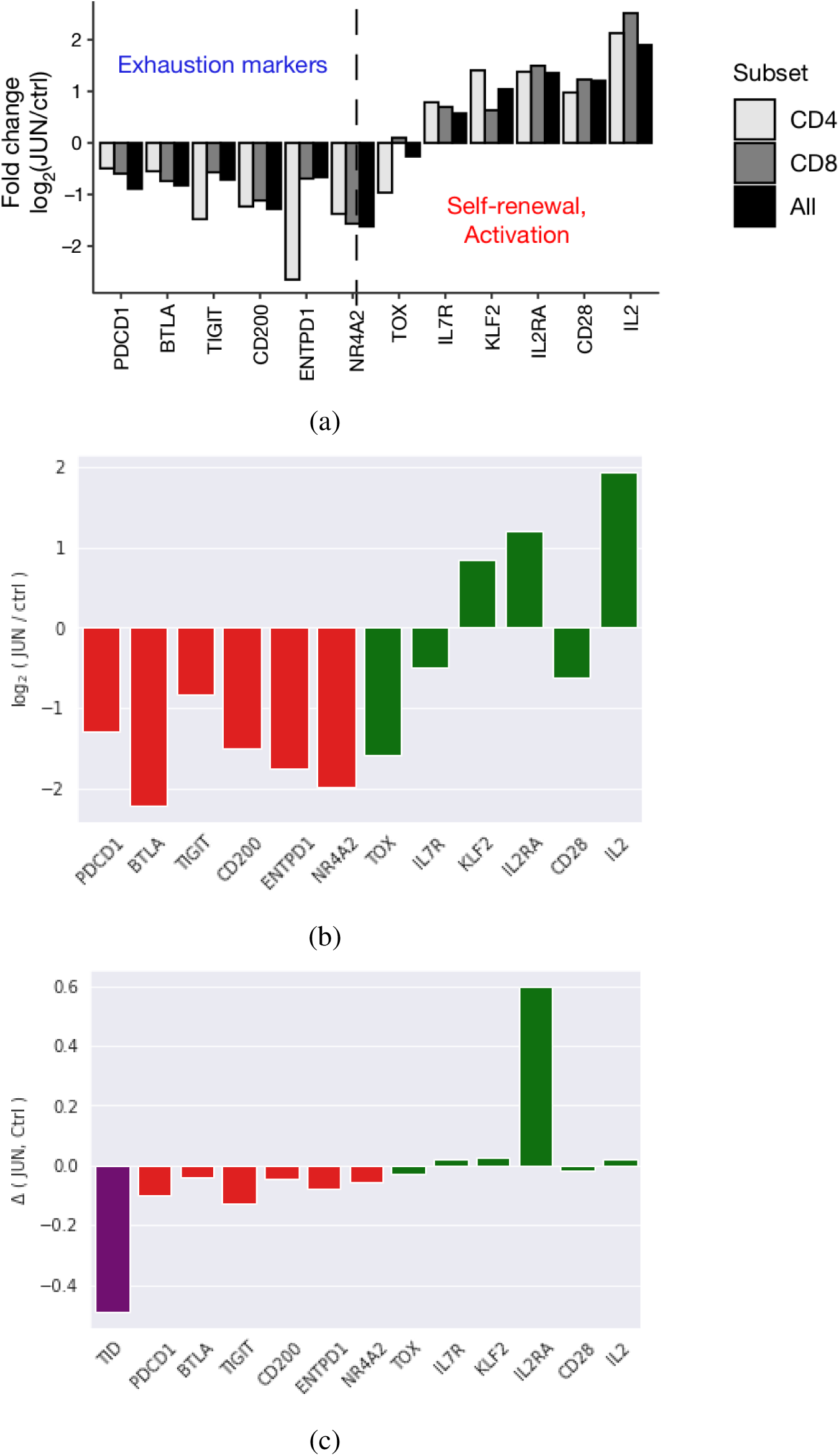
(a) Average comparisons between JUN and Control cells appearing in [5, figure 6g]. (b) Same comparisons using single-cell data. (c) Log-normalized comparisons, including TID.

**Figure 2:**
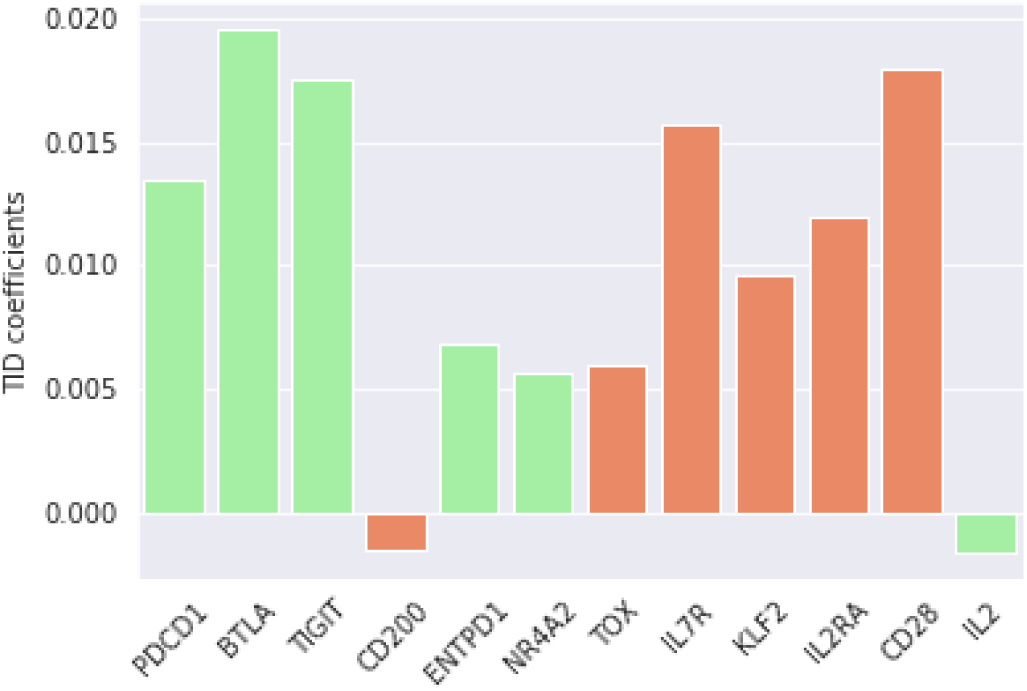
TID coefficients have “wrong” signs for many genes (in light red).

**Figure 3:**
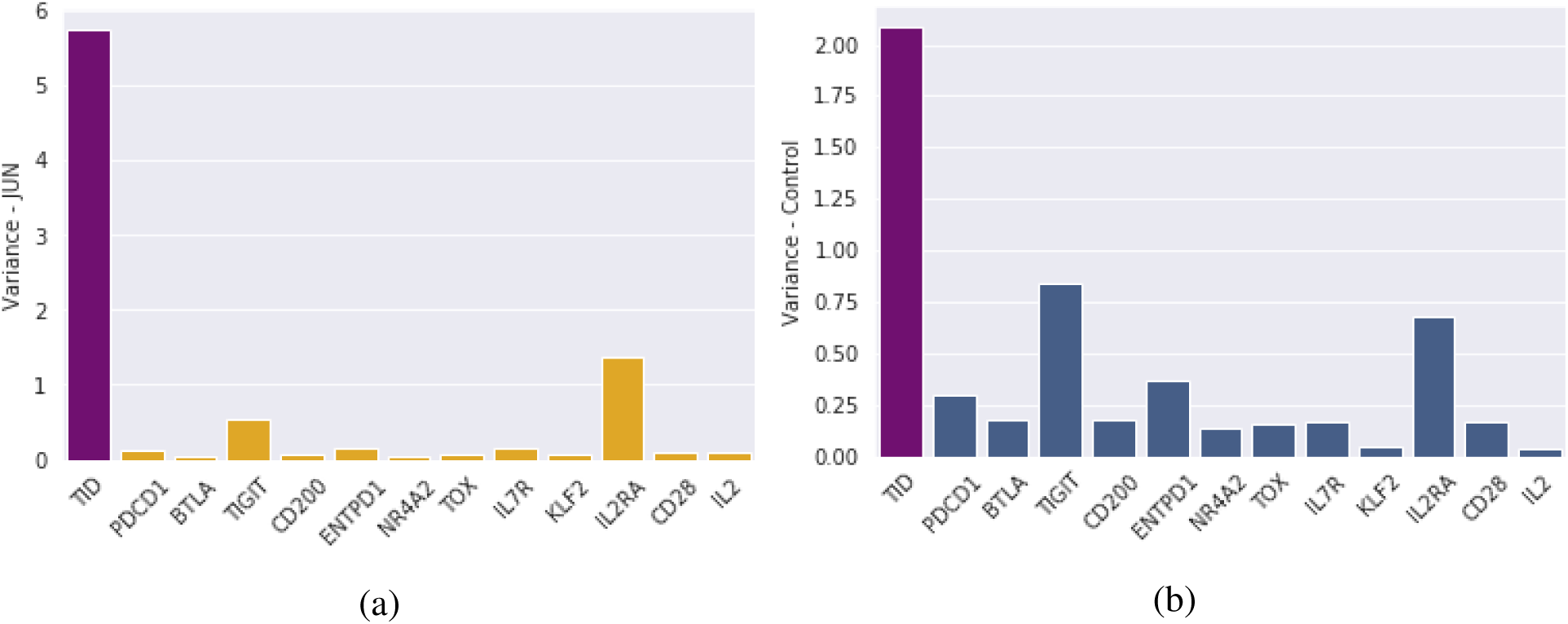
Variance for (a) JUN (b) Control CAR T cells.

**Figure 4:**
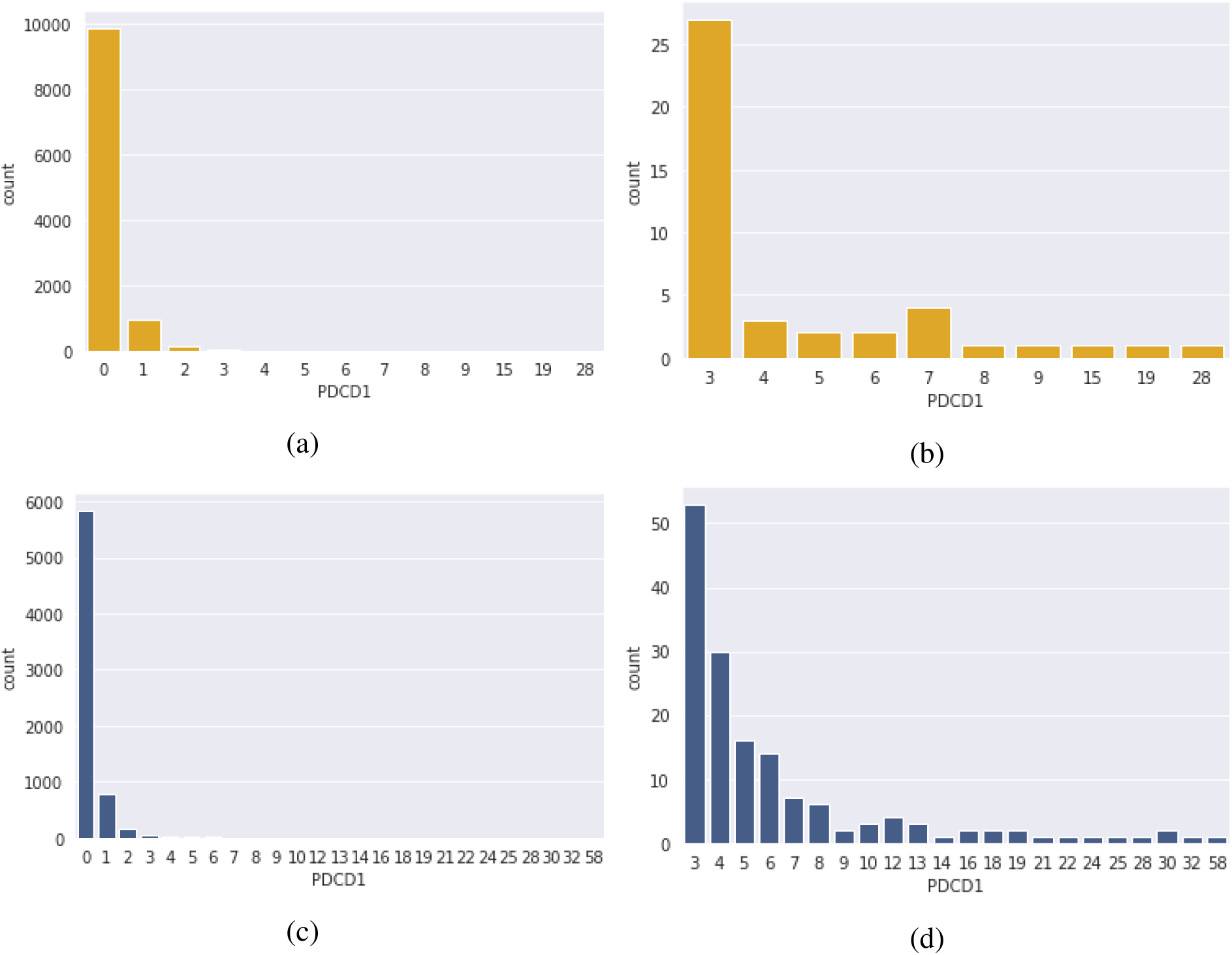
Counts of PDCD1 gene expression for (a) JUN CAR T cells (b) zoom for counts at least equal to 3 (c) Control CAR T cells (d) zoom for counts at least equal to 3.

**Figure 5:**
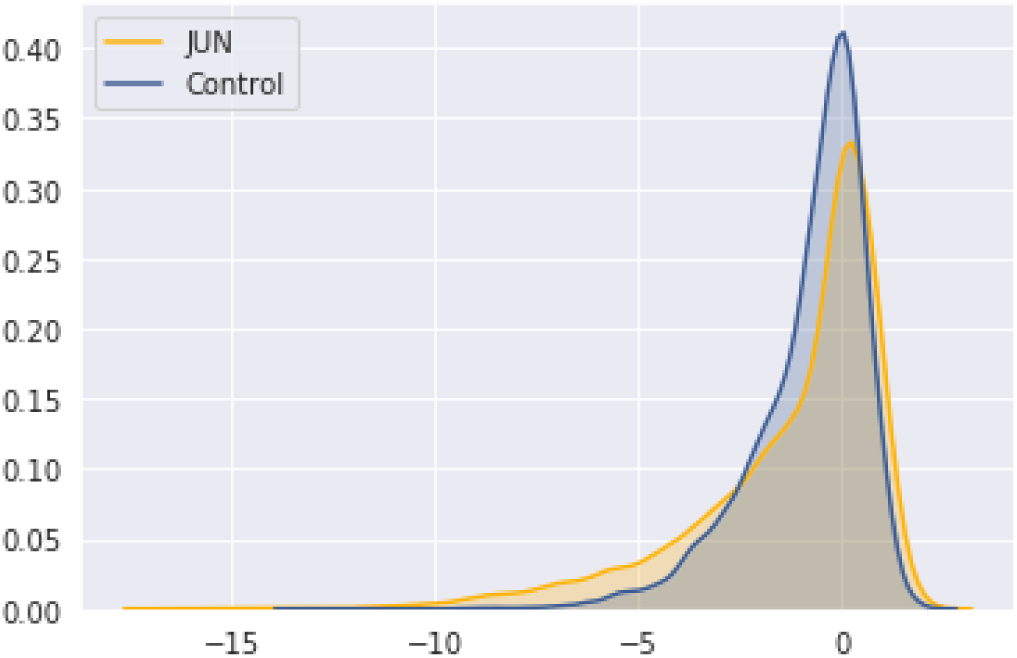
Distributions of TID scores for JUN and Control CAR T cells.

**Figure 6:**
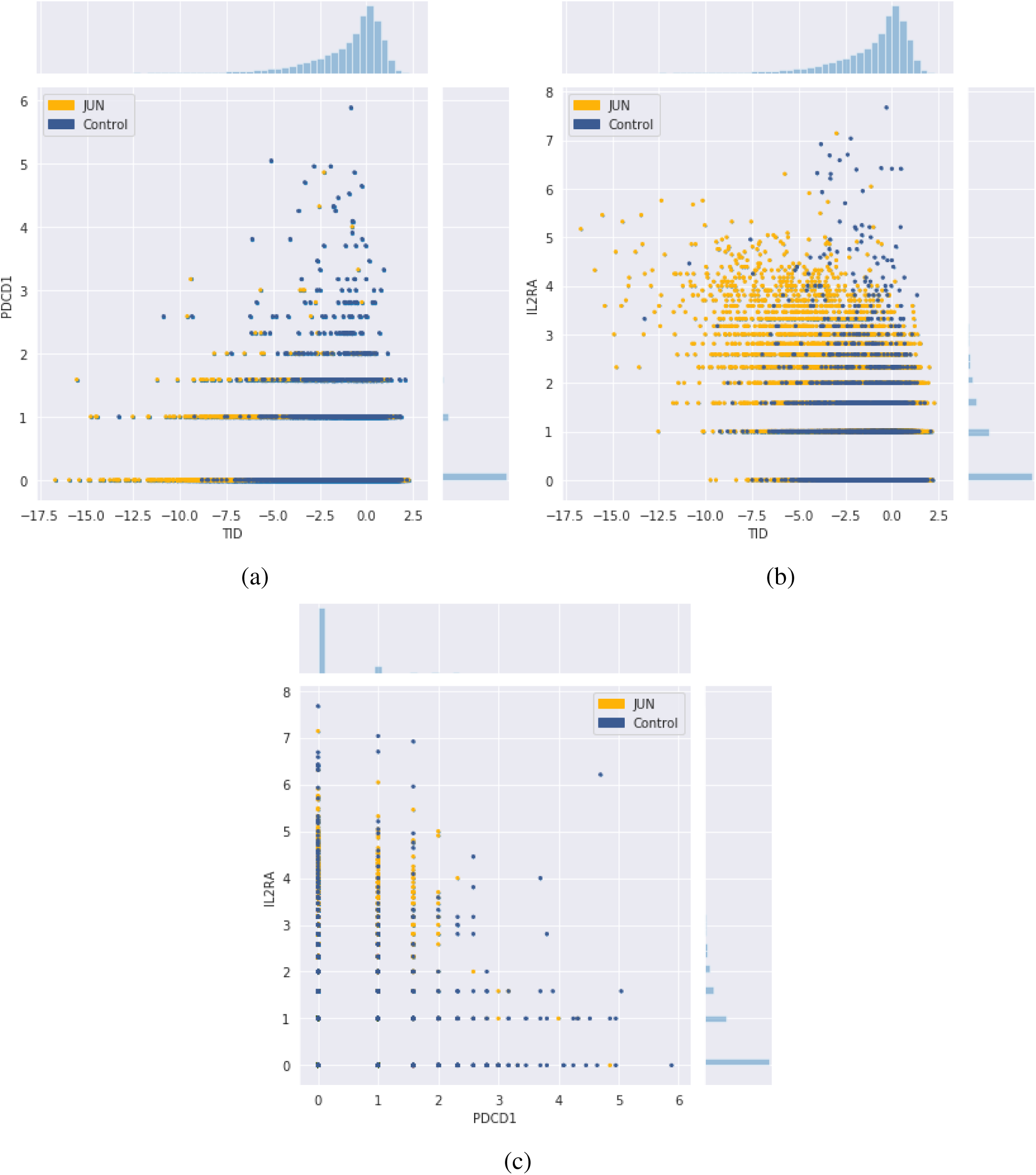
CAR T cells scores: (a) TID and PDCD1 (b) TID and IL2RA (c) PDCD1 and IL2RA.

- 6 signatures for exhaustion: PDCD1, BTLA, TIGIT, CD200, ENTPD1, and NR4A2;
- 6 signatures for activation and self-renewal: TOX, IL7R, KLF2, IL2RA, CD28, and IL2.

### TIDE and TID genome-wide signatures

TIDE is a genome-wide and bulk RNA-seq signature measuring Tumor Immune Dysfunction and Exclusion. TIDE combines dysfunction and exclusion signatures. Immune dysfunction signature integrates data from 73 human cancer studies, drawn from TCGA, PRECOG, and METABRIC databases, spanning various cancers such as melanoma, endometrial carcinoma, triple negative breast tumors, acute myeloid leukemia, and neuroblastoma [4, Supplementary table 2]. It is built by looking at correlations between cytotoxic T lymphocyte (CTL) level (estimated as the average gene expression level of CD8A, CD8B, GZMA, GZMB and PRF1), patient survival, and gene expression [4, figure 1b].

To keep the paper self-contained, we recall some details about how this genome-wide signature is obtained: TIDE uses the Cox Proportional-Hazards model to test how the interaction between a candidate gene *i*, of normalized expression level *h*_*i*_, and the CTL affects death hazard (estimated from survival):

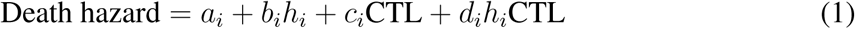

Coefficient *d*_*i*_ is the coupling coefficient between gene expression level *h*_*i*_ and T cell level CTL.

When *d*_*i*_ > 0, there is an antagonistic interaction between gene *i* and T cell level. A higher value of gene expression *h*_*i*_ will decrease the beneficial association between T cell level and survival (higher *h*_*i*_ will increase association between T cell level and death). For example, that’s the case for TGFB1.

When *d*_*i*_ < 0, that’s the opposite: there is a synergistic interaction between the gene *i* and T cell level. A higher value of gene expression *h*_*i*_ will increase the beneficial association between T cell level and survival. For example, that’s the case for SOX10.

When *d*_*i*_ = 0, there is no interaction between gene *i* and T cell level.

The resulting T cell dysfunction signature is a genome-wide vector, where the *z* score of each gene *i* is the interaction coefficient *d*_*i*_ divided by its standard error [4, Supplementary table 1] (that we still denote *d*_*i*_).

In our case, we further divide every *d*_*i*_ by the Euclidean norm of the vector (*d*_*i*_)_*i*∈Genome_ (a result that we still denote *d*_*i*_), so that we have:

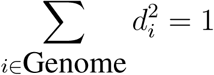

This last step makes homogeneity with single-gene signatures, which are “genome-wide” one-hot vectors (…, 0, 1, 0…) (i.e. 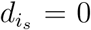 for single-gene signature *i*_*s*_, and *d*_*i*_ = 0 otherwise), and makes comparison possible. The resulting unit vector (*d*_*i*_) is what we define as the TID signature.

It’s important to note that TID is a bulk RNA-seq signature, which mixes all cells in the tumor micro-environment, not only T cells. For example, cancer cells can induce T cell dysfunction via SOX10 gene. TID is a signature about T cell dysfunction, not only exhaustion.

TIDE has been successfully applied to the prediction of outcomes of immune checkpoint inhibitor therapies, where it is one of the best classifiers [1]. However, TIDE could also be applied elsewhere. To our knowledge, that’s the first public attempt to apply TIDE to CAR T cell therapy for solid tumors.

## 1 TID is second-best at separating averages of JUN and control CAR T cells

### 1.1 Single-gene signatures averages

In [5, figure 6g] (copied here in figure 1a), there is a comparison of average expressions of 12 single-gene signatures. More precisely, [5, figure 6g] is a plot of:

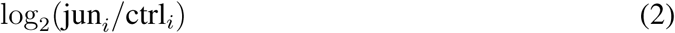

where jun_*i*_ and ctrl_*i*_ are the average expressions of gene *i* for JUN and Control CAR T cells respectively. They are given by:

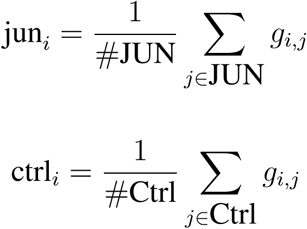

with #JUN and #Ctrl being the number of JUN and Control cells respectively, and with the nonnegative integer *g*_*i,j*_ being the expression of gene *i* for cell *j*.

[5, figure 6g] tells that, on average, JUN cells always express less exhaustion markers, and always more activation and self-renewal markers, than Control cells.

Based on single-cell gene expression data (publicly available in NCBI Gene Expression Omnibus, GEO series accession number: GSE136805), we tried to reproduce this [5, figure 6g]. Instead, we got figure 1b, which is different. We obtained relatively similar values for PDCD1, TIGIT, CD200, NR4A2, KLF2, ILR2A, and IL2. However, we obtained significantly different values for BTLA, ENTPD1, and even opposite signs for TOX, IL7R and CD28.

We have no explanation for this difference.

However, based on our figure 1b, the same conclusion remains, although a bit weakened, because compared with Control cells, JUN cells express all exhaustion markers less (6 over 6), and half of activation/self-renewal markers more (3 over 6).

### 1.2 TID average

In order to compare TID averages for JUN and control cells, we can’t generalize equation (2) to the genome-wide signature TID. That is to say, we can’t compute:

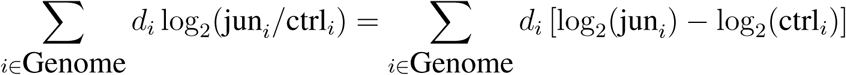

where *d*_*i*_ is the TID coefficient for gene *i*, because single-cell gene expression has a large amount of zeros. We have a lot of genes *i* such that: jun_*i*_ = 0 or ctrl_*i*_ = 0, with *d*_*i*_ ≠ 0. Therefore, we would take logarithms of zero and infinity.

Instead, in figure 1c, we plot, for TID:

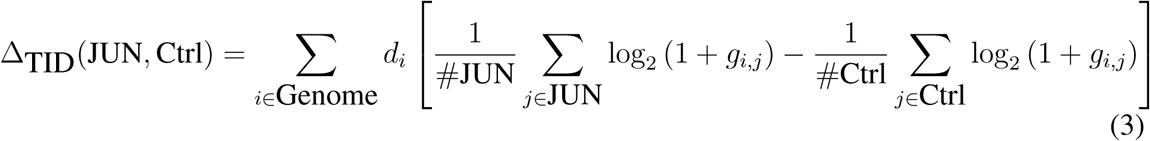

And for each of the 12 single-gene signatures *i*, we plot:

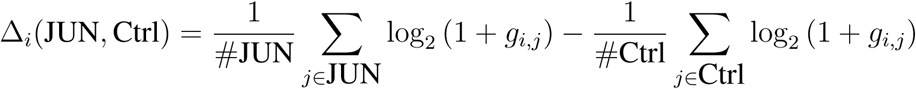

This corresponds to the best practice of *log-normalization*, i.e. applying the transformation *x* ↦ log_2_(1 + *x*) to gene expression. That is the recommended practice for TIDE. By the way, in TIDE definition in the hazard model of equation (1), the normalized gene expression is:

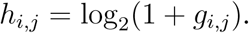

In figure 1c, we see that (log-normalized) TID average is lower in JUN than in Control cells. So according to TID, JUN cells are less dysfunctional than Control cells on average. Moreover, this difference is higher than with all other single-gene signatures, except IL2RA.

These results are even more remarkable when we see that *d*_*i*_ coefficients have the “wrong” sign for many genes: *d*_*i*_ is negative for the dysfunction gene CD200, and positive for most of activation genes: TOX, IL7R, KLF2, IL2RA, CD28 (figure 2).

However, contributions from those 6 genes with “wrong” signs together only weight 1.5 % of the average TID score.

One possible explanation for these “wrong” signs is that TID is not only a signature of T cell exhaustion, but a bulk RNA-seq signature of dysfunction, which takes into account different cells, such as other immune cells, cancer cells and stromal cells.

Another remark is that log-normalization is important for TID, because otherwise, the result is reversed: the average TID score for non-normalized JUN cells (−0.087) is larger than for Control cells (−1.148), which would mean that JUN cells would be more dysfonctional than Control cells.

Finally, in equation (3), TID average is a simplification that does not take into account the varying “CTL” weight of each T cell, which appears in equation (1) (i.e. the average gene expression level of CD8A, CD8B, GZMA, GZMB and PRF1): that’s a question to consider for adapting TIDE bulk signature to the single-cell setting.

## 2 CAR T cells heterogeneity is higher when measured with TID than with single-gene signatures, and is higher for JUN than for Control cells

Heterogeneity is an important feature of solid tumors, and a factor of therapy resistance [3]. Single cell RNA-seq facilitates analyzing this heterogeneity. With data available, we can study CAR T cells heterogeneity within JUN and Control cells categories. [5] measures CAR T cells heterogeneity in different ways, in [5, figure 6] and [5, extended figure 10]. However, they didn’t measure variance between cells.

The variance of gene *i* log-normalized expression, for JUN cells, is given by:

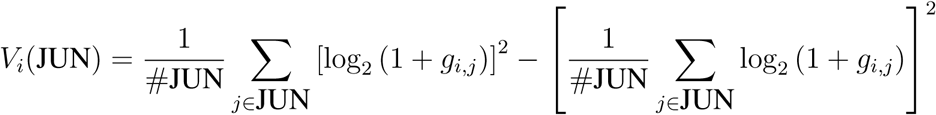

where the nonnegative integer *g*_*i,j*_ is the expression of gene *i* for cell *j*.

The TID variance for JUN cells is given by:

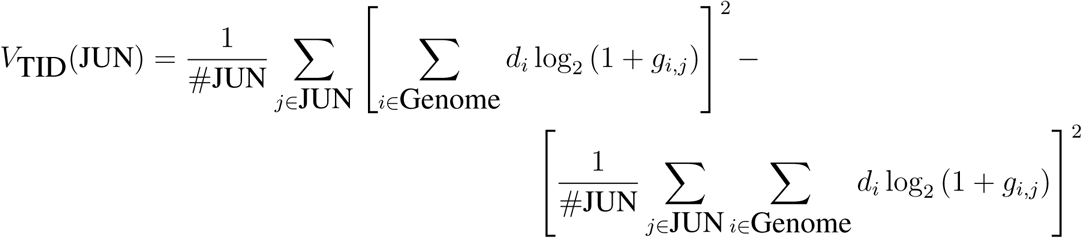

We did not need to normalize gene expression by substracting the average expression, because variance does not change by subtracting the average.

The variance for Control cells is given by similar formulas.

In figures 3a and 3b, we can see that TID variance is the highest, by a large margin. Moreover, TID variance is higher in JUN than in Control cells.

This suggests that there is heterogeneity in JUN cells dysfunctional states. This possibility could have been overlooked by only considering single-gene signatures. This heterogeneity could potentially provoke therapy failures at later stages in drug development.

In any case, this hypothesis should be tested in more precise mouse experiments, with longer follow-ups.

Indeed, survival data in [5, extended figure 9h] is very limited. It only shows 50 days follow-up. Up to 50 days, all mice engrafted with solid tumors, and then treated with JUN-Her2-BBz CAR T, survive. On the other hand, all mice treated with Control Her2-BBz CAR T die between 28 days and 38 days. Therefore, survival data only shows that JUN-Her2-BBz CAR T treatment extends mice life by 22 days. There is no update about what happens after, despite that [5] manuscript was submitted in September 2018.

Concern about short follow-up in [5, extended figure 9h] is amplified by information shown in [5, extended figure 9e]. In this experiment, all mice injected with liquid tumors, and then treated with JUN-CD19-BBz CAR T, die between 28 days and 48 days. On the other hand, all mice treated with Control CD19-BBz CAR T die between 19 and 24 days. So in this case, JUN CAR T engineering extends mice life by 30 days at most.

Another possibility is that these high variances might not reflect a real-world phenomenon, and only come from noise in the TID signature, which has a lot of counter-intuitive coefficients, as it appears in figure 2. This question would require further investigations with different single-cell datasets.

## 3 TID and single-gene signatures distributions don’t show inter-mouse heterogeneity

To refine our observations about variance, we plot distributions.

In particular, these plots might be a way to detect inter-mouse heterogeneity. In single-cell data published by [5], mouse labels are missing: only their categories (JUN or Control) are provided. Moreover, mice survival follow-up is too short to be informative about heterogeneity in JUN mice.

That’s why workarounds to detect inter-mouse variation can be useful, and multi-modal distributions could be a hint. They are superpositions of uni-modal (single-peaked) distributions. However, it is still possible to have uni-modal distribution with inter-mouse variation.

This is an important question because without inter-mouse variation, it would mean that TID variance in CAR T cells observed in section 2 is intra-mouse and intra-tumoral.

Moreover, immuno-oncology is often about personnalized oncology. In the case of immune checkpoint inhibitors, there is large inter-patient heterogeneity in clinical outcomes.

### 3.1 Single-gene signatures

Single-gene signatures take integer values, so it’s convenient to plot histograms. We did it for PDCD1, in figures 4. We didn’t log-normalized gene expression, for better readability.

We see a sharp and almost regular decrease in cell numbers, when gene expression per cell grows. As expected, the decrease is sharper for JUN than for Control cells. We don’t see any significantly multi-modal distribution, although this does not rule out inter-mouse variation, which appears in [5, extended figure 10b] for PD-1.

Other single-gene signatures are left as exercises. Another exercise is to check whether the distribution is geometric.

### 3.2 TID signature

TID signature is a linear combination of thousands of single-gene signatures, and therefore, by approximation, it takes continuous values. In figure 5, we plot distribution densities of TID scores, for JUN and Control CAR T cells. Both distributions are skewed and unimodal (single-peaked), with JUN cells distribution having a thicker tail, hence having lower mean and higher variance than Control cells distribution.

Again, we don’t see any multi-modal distribution that could suggest significant inter-mouse variation, although this possibility is not ruled out either.

An exercise is to check whether distributions are log-normal, and compute their parameters.

## 4 TID has a one-sided correlation with single-gene signatures

We examine the relationship between TID signature and single-gene signatures of a CAR T cell. In figure 6a, we plot TID scores against PDCD1 scores, for both JUN and Control CAR T cells (all gene expressions are log-normalized). Both are T cell dysfunction signatures.

We observe that low PDCD1 cells can have either low or high TID levels. Likewise, high TID cells can either have low or high PD1. However, low TID cells never have high PDCD1 levels. Moreover, JUN cells more often have low TID, low PDCD1 scores, and Control cells more often have high TID, high PDCD1 scores. That’s consistent with the finding that JUN are less dysfunctional than Control cells.

In contrast, in figure 6b, we plot TID scores against IL2RA, which is an activity signature.

Cells with low levels of TID can have high IL2RA, but can’t have low IL2RA.

In figure 6c, we compare two single-gene signatures, PDCD1 and IL2RA. We make analogous observations, and in particular, high PDCD1 cells almost never have high IL2RA.

These partial correlations show that TID is complementary with single-gene signatures.

## 5 CAR T cells with low TID levels make distinct clusters

As in [5, extended figure 10f and 10g], we plot 2-dimensional UMAP projections of CAR T cells. This gives a qualitative way to visualize similarity between cells. UMAP is an embedding of high dimensional data, analogous to t-SNE, with improvements in terms of computational speed and global distance preservation [6].

In figure 7b, we project expression data. We obtain a different result from [5, extended figure 10f] (reproduced in figure 7a). This difference might be due to UMAP stochastic nature [6, section 5.2], which gives different results from run to run, like t-SNE.

**Figure 7:**
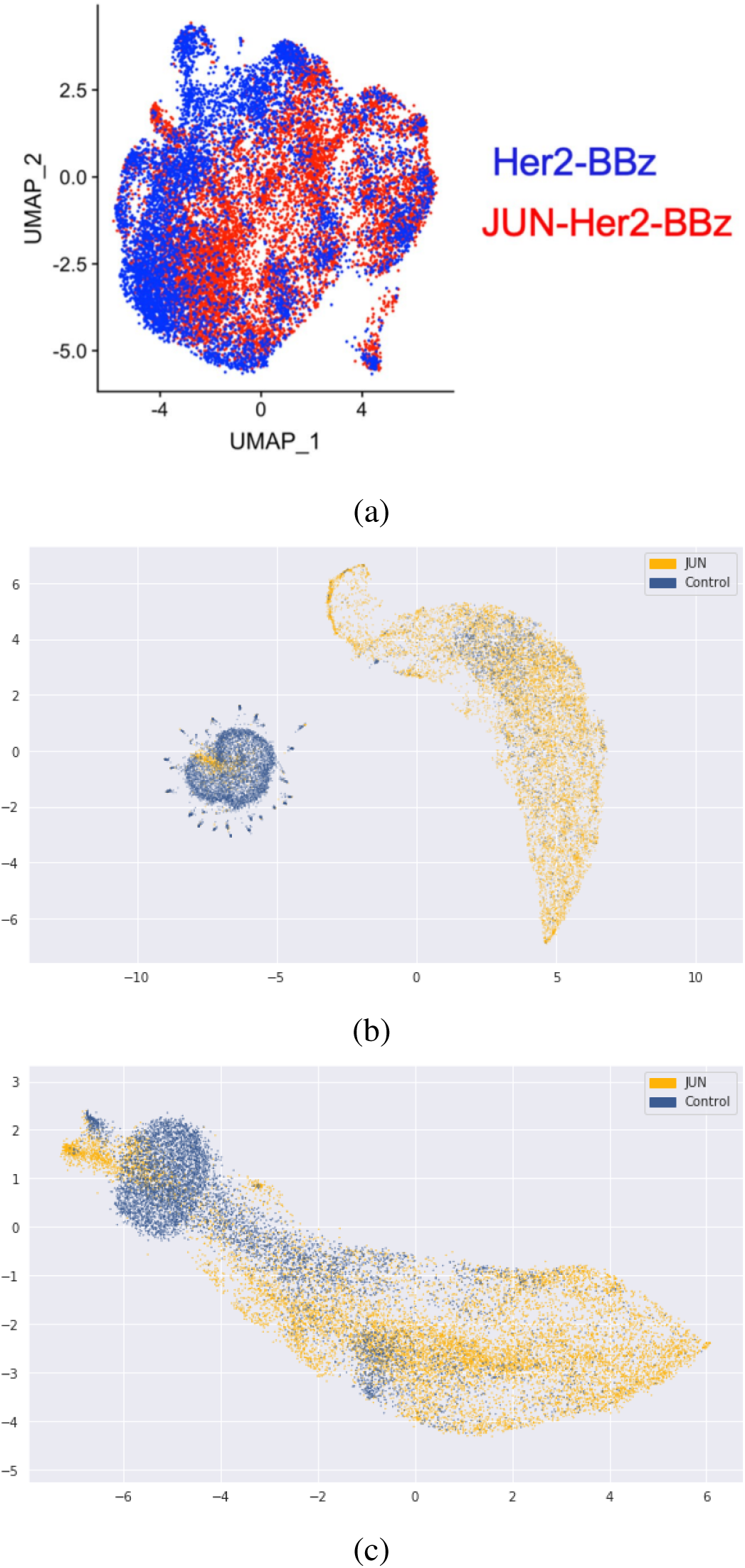
UMAP visualizations of CAR T cells (a) as it originally appears in [5, extended figure 10f] (JUN cells: JUN-Her2-BBz, Control cells: Her2-BBz), (b) as we reproduce it (c) with log-normalized data.

In figure 7c, we project the log-normalized gene expression data, which is more relevant in our study. In both cases (with and without log-normalization), we can see that most of JUN and Control cells belong to different clusters. This confirms that JUN cell engineering significantly impacts cell activity (this was less obvious from UMAP visualization in [5, extended figure 10f]).

Moreover, the JUN cluster is larger than the control cluster, which supports our conclusion drawn from variance computation in section 2, according to which JUN cells are more heterogeneous than Control cells.

In figure 8a, we can see that zones of lowest TID scores for JUN cells (the least TID-dysfunctional JUN cells) are located at the tip, the most further away from Control cells. For Control cells, zones of lowest TID scores are the most further away from the core cluster.

**Figure 8:**
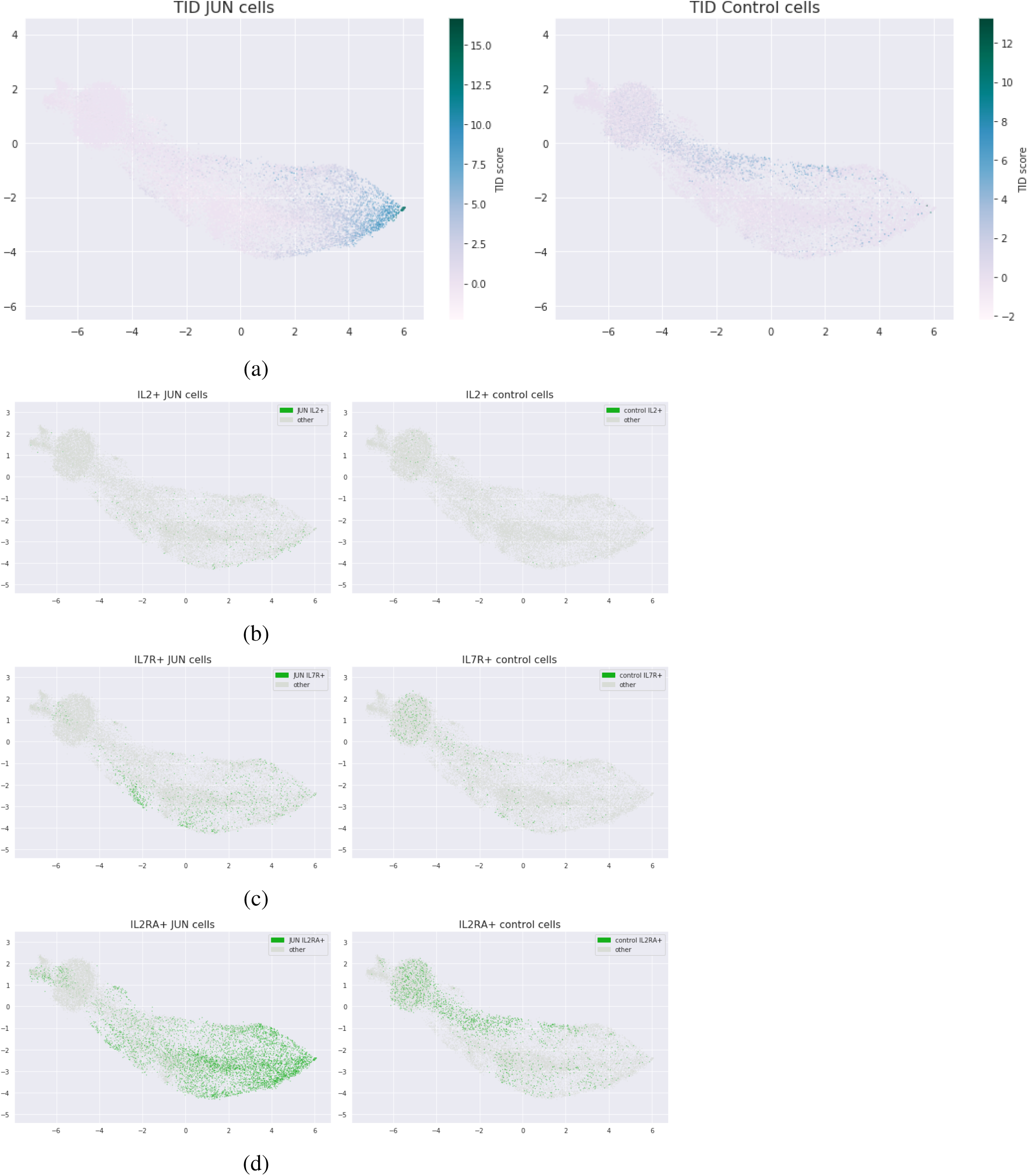
UMAP visualizations of CAR T cells coloring (a) TID scores (continuous scale) (b) IL2 > 0 (c) IL7R > 0 (d) IL2RA > 0

Figure 8b shows cells expressing IL2. They overlap with TID-low cells but they can be located elsewhere too.

The same conclusion holds for cells expressing IL7R (figure 8c): they overlap with the TID-low cluster, but most of their concentration is located far away.

Figure 8d shows that cells expressing IL2RA are spread out.

Other single-gene signatures are left as an exercise.

In conclusion, UMAP visualizations show that the least TID-dysfunctional CAR T cells form specific sub-clusters.

## Conclusion

We have shown how genome-wide signature TID can enrich analysis of CAR T for solid tumors scRNA-seq data. First, it confirms that on average, JUN CAR T cells are less dysfunctional than Control CAR T cells. However, it also suggests to be careful about JUN CAR T cells heterogeneity, which brings uncertainty. In conclusion, TID helps de-risking this difficult pipeline in oncology drug development.

## Acknowledgement

I would like to thank Peng Jiang for correspondence.

## Data and Code Availability

Data and code (including a hands-on tutorial) can be available upon request at: https://melwy.com

